# Gold Nanocages with a Long SPR Peak Wavelength as Contrast Agents for Optical Coherence Tomography Imaging at 1060 nm

**DOI:** 10.1101/2025.03.09.642209

**Authors:** Yongping Chen, Jiefeng Xi, Du Lee, Jessica Ramella-Roman, Xingde Li

## Abstract

There has been growing interest in optical coherence tomography (OCT) imaging at a wavelength of 1060 nm. However, potential contrast agents for OCT imaging at this specific wavelength has not been thoroughly investigated. In this study, we present the synthesis and optical characterization of gold nanocages with a small edge length (∼65 nm) and a surface plasmon resonance peak around 1060 nm. These nanocages represent a class of potential contrast agents for OCT at 1060 nm. OCT imaging experiments were conducted on phantoms and *in vivo* mouse tissues using a 1060-nm swept-source OCT system, demonstrating significant enhancement in imaging contrast due to the presence of the gold nanocages.

**GRAPHICAL ABSTRACT:** Gold nanocages with a long SPR peak wavelength as OCT imaging contrast agents at 1060 nm

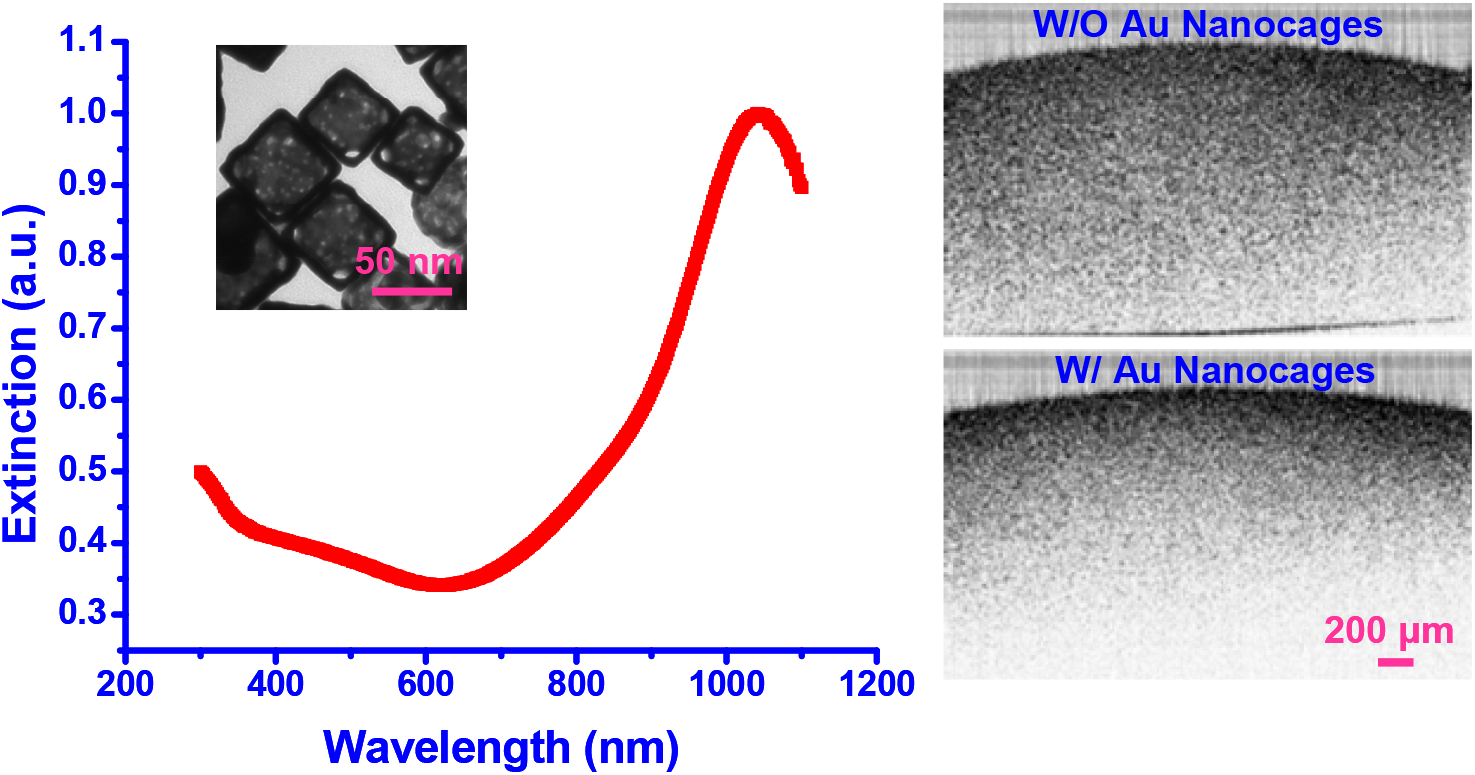

## INTRODUCTION

Optical Coherence Tomography (OCT) is a noninvasive imaging technology that enables real-time, high-resolution visualization of biological tissues [1]. Swept-source OCT (SS-OCT) has emerged as a powerful advancement, offering fast a-line scan rates and high detection sensitivity. Conventional OCT imaging contrast is primarily governed by the intrinsic optical scattering and absorption properties of biological tissues (often dominated by scattering). Similar to other imaging modalities such as X-ray computed tomography, ultrasound, and magnetic resonance imaging, the use of exogenous contrast agents can significantly enhance the imaging capabilities of OCT and more importantly molecular specificity. Contrast enhancement has been achieved through the use of various agents, including dyes [2], microspheres [3], microbubbles [4], and gold nanoparticles, such as nanorods [5,6], nanoshells [7,8], and nanocages [9,10].

Gold nanoparticles are particularly attractive for OCT contrast enhancement due to their bioinertness, customizable absorption and scattering cross-sections, and tunable surface plasmon resonance (SPR) peaks, which can be matched to specific OCT light sources. Among gold nanoparticles, gold nanocages—structured particles with a hollow interior and a thin yet robust porous wall—have garnered attention. These particles are synthesized using a galvanic replacement reaction between Ag nanocubes and HAuCl_4_ in an aqueous solution. The SPR peaks of gold nanocages can be tuned to the near-infrared (NIR) region by adjusting the thickness and porosity of their walls [11]. Compared to gold nanorods and nanoshells, gold nanocages exhibit stronger absorption in the NIR region while maintaining a smaller size, which is crucial for effective delivery through tissues. This makes gold nanocages particularly well-suited as contrast agents for OCT imaging. Our previous work demonstrated that gold nanocages enhanced OCT intensity and spectroscopic imaging contrast at wavelengths around 800 nm [9,10].

OCT imaging using light sources centered around 1060 nm has become very popular, as this wavelength offers deeper tissue penetration due to lower optical scattering and minimal dispersion compared with OCT at 800 nm, and lower absorption when compared with OCT at 1300 nm [12,13]. Recent studies have explored the use of polypyrrole nanoparticles as contrast agents for OCT imaging at 1060 nm [14]. However, most gold nanoparticles have SPR wavelengths below 1000 nm, making them unsuitable as contrast agents for OCT imaging at this wavelength. In this study, we investigate the use of gold nanocages with an SPR peak around 1060 nm to enhance OCT imaging contrast.

While gold nanocages with SPR peaks across a broad wavelength range can be synthesized following established protocols [11], it remains challenging to produce small gold nanocages with SPR peaks at longer wavelengths (i.e., beyond 950 nm). This challenge arises because the nanocages tend to collapse during the final stage of the galvanic replacement reaction, particularly when the concentration of HAuCl_4_ increases. In this work, we demonstrate that gold nanocages with SPR peaks shifting to approximately 1062 nm and beyond can be synthesized by reducing the reaction temperature and titration rate by 3-4-fold. We also quantitatively characterize the optical properties of these nanocages using an integrating sphere and the inverse adding-doubling method [15]. Finally, we present real-time OCT imaging with a 1060 nm swept source, performed on both tissue phantoms and *in vivo* mouse tissues, illustrating the contrast enhancement achieved with the gold nanocages.

## METHODS

### Synthesis of gold nanocages with an SPR wavelength above 1000 nm

Gold nanocages with a longer SPR wavelength were synthesized following the galvanic replacement reaction principle as described by Skrabalak et al. [11], with a modified procedure. Briefly, 100 µL of an aqueous silver nanocube solution at a concentration of approximately 6 nM was first dispersed in 5 mL of deionized water containing poly(vinyl pyrrolidone) (1 mg/mL) in a 50 mL flask under magnetic stirring at room temperature (instead of the conventional ∼90-100 °C) for approximately 10 minutes. Then, an aqueous solution of HAuCl_4_ (0.1 mM) was added to the flask using a two-channel syringe pump at a rate of 0.25 mL/min (about one-third of the conventional titration rate) while maintaining magnetic stirring. During the titration of the HAuCl_4_ solution, a series of color changes were observed, and the position of the SPR peaks of the gold nanocages was monitored using a UV-Vis-NIR spectrophotometer. The titration of the HAuCl_4_ solution was stopped once an appropriate SPR peak was reached. The reaction solution was then transferred to a 50 mL centrifuge tube and centrifuged at 5000 rpm for approximately 5 minutes. The supernatant was discarded, and the gold nanocages were redispersed in a saturated NaCl solution to dissolve and remove AgCl, which was generated during the galvanic replacement reaction. Finally, the solution was centrifuged at 10,000 rpm for about 10 minutes. The gold nanocages were thoroughly washed with deionized water and redispersed in 18.1 MΩ·cm E-pure water for further use.

### Measurement of optical properties of gold nanocages

The optical properties of the gold nanocages were quantitatively characterized using the well-established integrating sphere method in conjunction with a spectrophotometer (Ocean Optics, Dundee, FL). Measurements were conducted over the wavelength range of 1000 to 1200 nm. The nanocage solutions (∼1 nM) in glass vials were ultrasonicated in a water bath for 20 minutes prior to measurement. Custom-made glass cuvettes with a thickness of 1 mm and dimensions of 25.5 × 25.5 mm were used to contain the solutions during the integrating sphere measurements.

### Gold nanocages as contrast agents for OCT imaging at 1060 nm

Swept-source OCT (SS-OCT) imaging was performed on phantoms and mouse tissues *in vivo*, both with and without the administration of gold nanocages. The as-synthesized gold nanocages had an edge length of approximately 65 nm, an SPR peak around 1040 nm, and a full-width-at-half-maximum (FWHM) spectrum bandwidth of ∼300 nm, which overlaps with the 1060-nm OCT source spectrum. For phantom experiments, each phantom was prepared from 5% gelatin embedded with 0.1% (by weight) Titania granules to mimic background tissue scattering. The scattering coefficient (µs) of the phantom was estimated to be approximately 5/mm, based on the decay curve of the OCT image. Gold nanocages were added to the phantom to a final concentration of 0.5 nM (for phantom experiments, the concentration of gold nanocages could be varied; 0.1 nM also works for OCT imaging). SS-OCT imaging was conducted using a swept laser with a center wavelength of 1060 nm and a FWHM spectral bandwidth of ∼65 nm.

For *in vivo* tissue experiments, a 50 µL aqueous solution of gold nanocages at a concentration of approximately 1 nM was locally injected into the tail tissue of an NCR nude mouse (with the concentration of nanocages remaining at about 1 nM in the tissue) under anesthesia. OCT imaging was performed using the same method as described above. Animal experiments in this study were approved by the Institutional Animal Care and Use Committees at Johns Hopkins University.

## RESULTS

### Gold nanocage synthesis, influence of synthesis conditions, and characterization

Figure 1 shows the UV-vis-NIR extinction spectra of the gold nanocages versus the titrated volume of the HAuCl_4_ solution to the reaction solution. The SPR peak continuously shifted towards a longer wavelength as the total volume of HAuCl_4_ increased, and an SPR peak wavelength of around 1062 was achieved after 5.2 mL of HAuCl_4_ solution was titrated. Figure 2 illustrates representative transmission electron microscopy (TEM) images of the nanostructures that were collected at different stages during the synthesis reaction. Figure 2A shows the silver nanocubes of an ∼60 nm edge length which served as the templates for gold nanocage synthesis.

**Figure 1.**
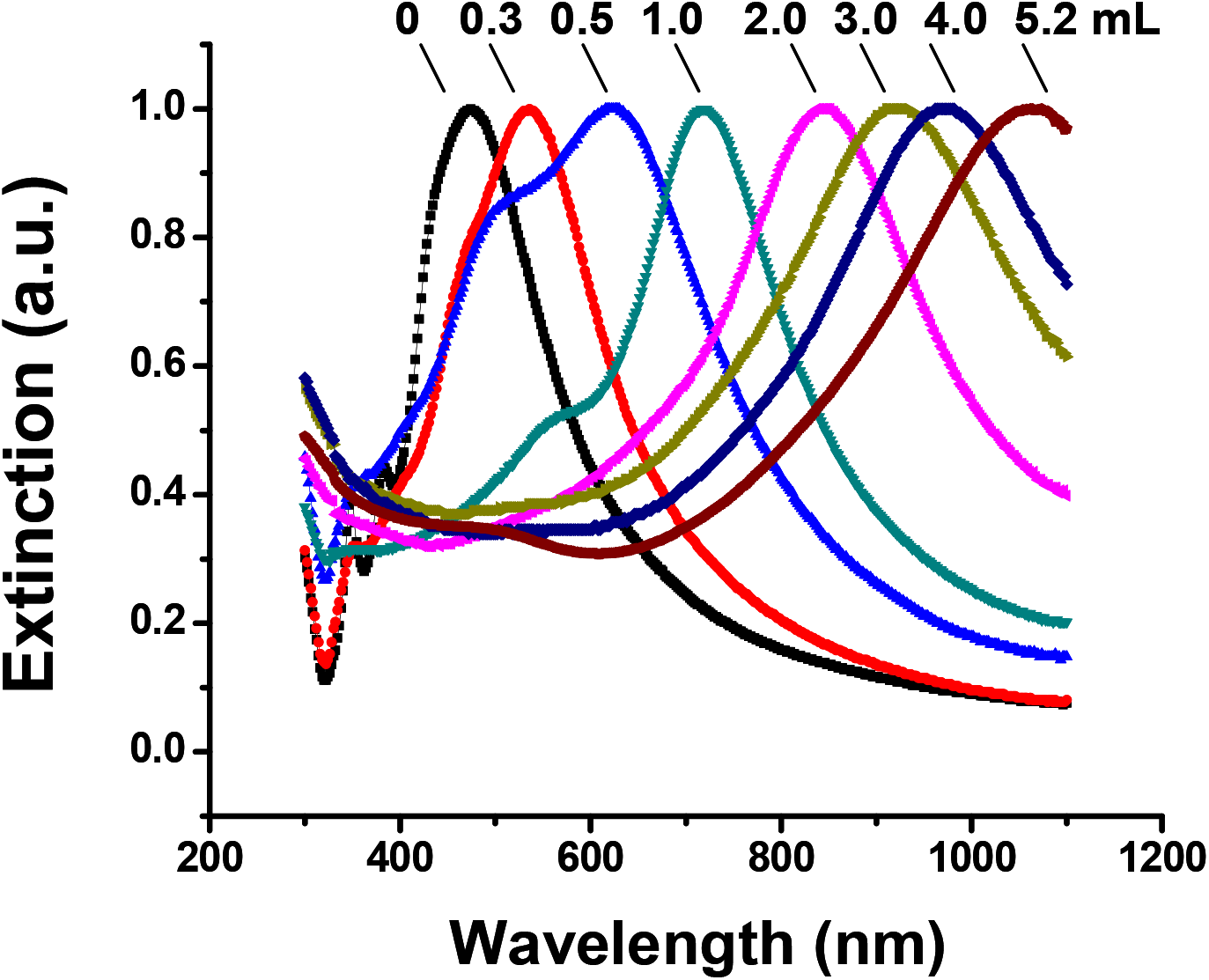
UV-vis-NIR spectra of the samples obtained by reacting silver nanocubes with different volumes of 0.1 mM HAuCl_4_ solution under room temperature with a titration rate of ∼0.25 mL/min. The SPR peak of the final product - gold nanocages - can be conveniently tuned to ∼1062 nm after 5.2 mL of HAuCl_4_ solution was titrated.

**Figure 2.**
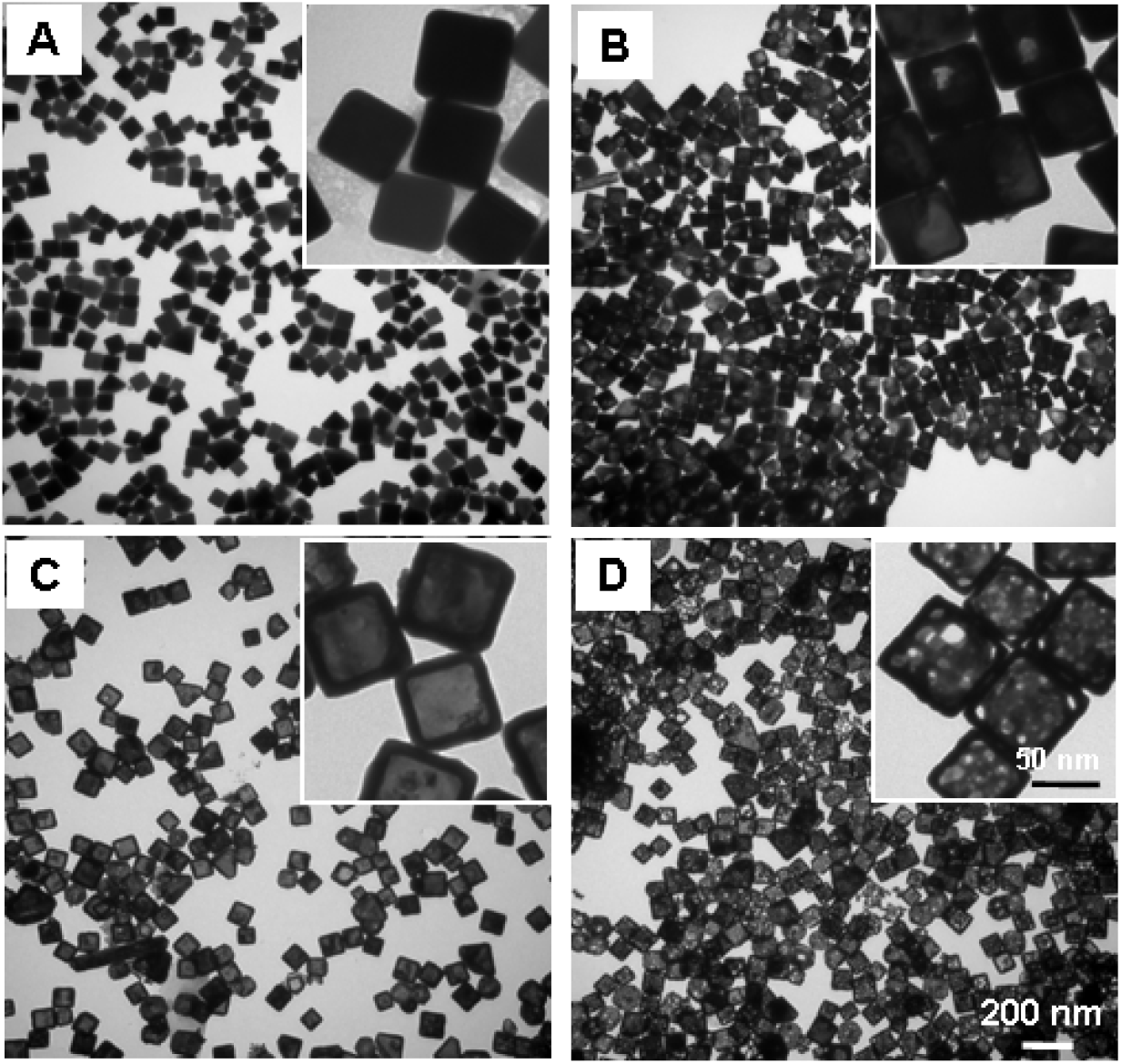
TEM images of (A) 60-nm Ag nanocubes which serve as the templates for gold nanocage synthesis; (B) Au/Ag alloy nanostructures with a pinhole observed on the wall during the early stage of the galvanic replacement reaction between the Ag nanocubes and HAuCl_4_ in aqueous solution; (C) Au/Ag alloy nanoboxes with an SPR peak around 782 nm; and (D) Au nanocages (after continuous dealloying) as the end product of the galvanic replacement reaction with a longer SPR peak around 1062 nm. Pores could be clearly observed on the wall of the nanocages. The insets show a zoomed-in TEM image of the Ag nanocubes, Ag/Au nanostructures, nanoboxes, and Au nanocages, respectively.

After Ag nanocubes reacted with HAuCl_4_ in solution, a pinhole could be observed on the wall (Fig. 2B); the SPR peak at this early reaction stage was around 622 nm. As more HAuCl_4_ solution was added, the interior of the Ag nanocube templates was continuously dissolved to yield a hollow nanobox through a combination of galvanic replacement and alloying between Ag and Au (Fig. 2C), and the SPR peak continuously shifted to a longer wavelength (e.g., around 782 nm). At this stage, the wall thickness of the nanocages measured from the TEM images was about 10.7 ± 0.6 nm. In the later stage, gold nanocages, with a hollow interior and porous wall (see Fig. 2D), were obtained through dealloying of the nanoboxes’ walls ^11,16^ and the SPR peak reached a much longer wavelength (e.g., around 1062 nm). At this stage, the measured wall thickness was about 7.2 ± 0.4 nm.

To demonstrate the effect of the reaction temperature and titration rate, conventional synthesis conditions were used, i.e., at a temperature of ∼ 90-100 °C and a HAuCl_4_ solution titration rate of 0.75 mL/min, while the rest of the reaction conditions remained the same. Under these conditions, dealloying caused the nanocages to collapse and form small gold fragments or clusters of irregular shapes, resulting in a second SPR peak around 550 nm when trying to shift the SPR peak beyond 930 nm. Figures 3A and 3C show a UV-visible-NIR extinction spectrum and a TEM image of the collapsed gold nanocages, respectively. When we decreased the reaction temperature to room temperature and the titration rate to 0.25 mL/min, the reaction between silver nanocubes and HAuCl_4_ was found to be stable, resulting in gold nanocages with SPR peaks above 1000 nm. Figures 3B and 3D respectively show a representative extinction spectrum and a TEM image of gold nanocages with an SPR peak at 1062 nm which were synthesized under the modified conditions. From the TEM image the average edge length of the nanocages was found to be 65.2 ± 4.6 nm.

**Figure 3.**
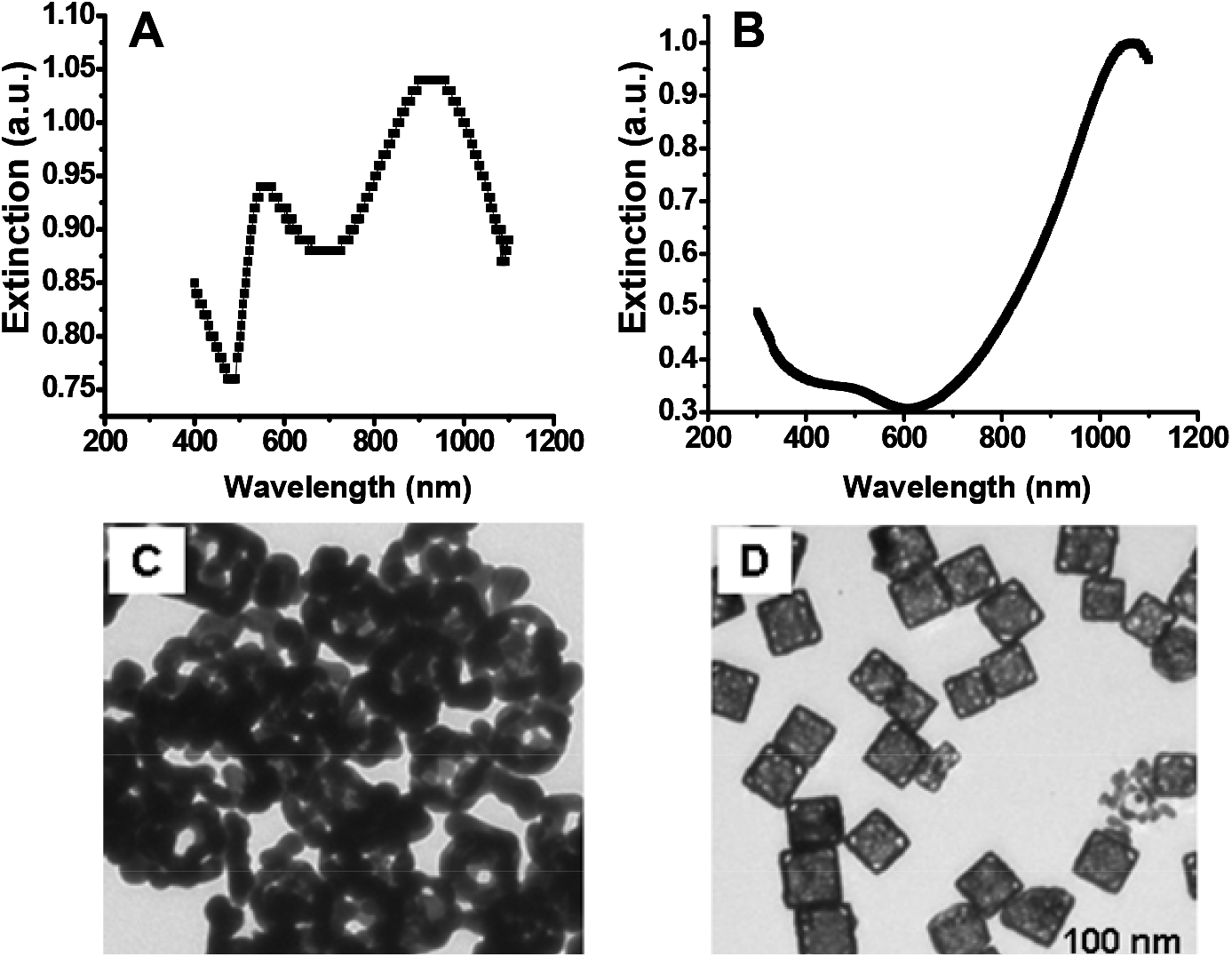
UV-vis-NIR extinction spectra (top) and TEM images (bottom) of gold nanocages. (A and C): under the conventional synthesis conditions (i.e., ∼ 90-100 °C with a titration rate of ∼0.75 mL/min), the reaction resulted in an SPR peak around 930 nm and a new peak around 550 nm. (B and D): under the modified synthesis conditions (i.e., room temperature with a reduced titration rate of ∼0.25 mL/min), the reaction resulted in a single SPR peak around 1062 nm.

The optical properties (i.e., the absorption and reduced scattering coefficients) of the gold nanocages were measured using the integrating sphere method. According to discrete dipole approximation (DDA) calculations, the anisotropy factor (g) is about zero for gold nanocages; hence the scattering coefficient (µ_s_) is the same as the measured reduced scattering coefficient [µ_s_’ = (1-g) µ_s_ = µ_s_]. The absorption (σ_abs_) and scatting (σ_sca_) cross-sections can then be obtained from the absorption (µ_a_) and scattering (µ_s_) coefficient according to σ_abs, sca_ = µ_a, s_ / ([C] *N_A_), where [C] is the concentration of gold nanocages and N_A_ is the Avogadro constant. The absorption (σ_abs_) and scattering (σ_sca_) cross-sections of a gold nanocage at 1060 nm, were found to be (1.55 ± 0.38) x 10^−15^ m^2^ and (3.14 ± 0.09) x 10^−16^ m^2^, respectively, from which the ratio of σ_abs_ to σ_sca_ is calculated to about 4.9, suggesting the interaction of light with these nanocages are dominated by optical absorption.

### Gold nanocages as contrast agents for OCT imaging at 1060 nm

Intensity-based SS-OCT imaging was performed on phantoms and mouse tail tissues *in vivo*, both with and without the administration of gold nanocages. Figures 4A and 4B show intensity-based SS-OCT images of the phantom without and with nanocages, respectively. The representative decay curves of backscattering intensity along imaging depth for both cases are compared in Fig. 4C. The experimental results show that the presence of gold nanocages significantly increased the overall attenuation (dominated by absorption increase).

**Figure 4.**
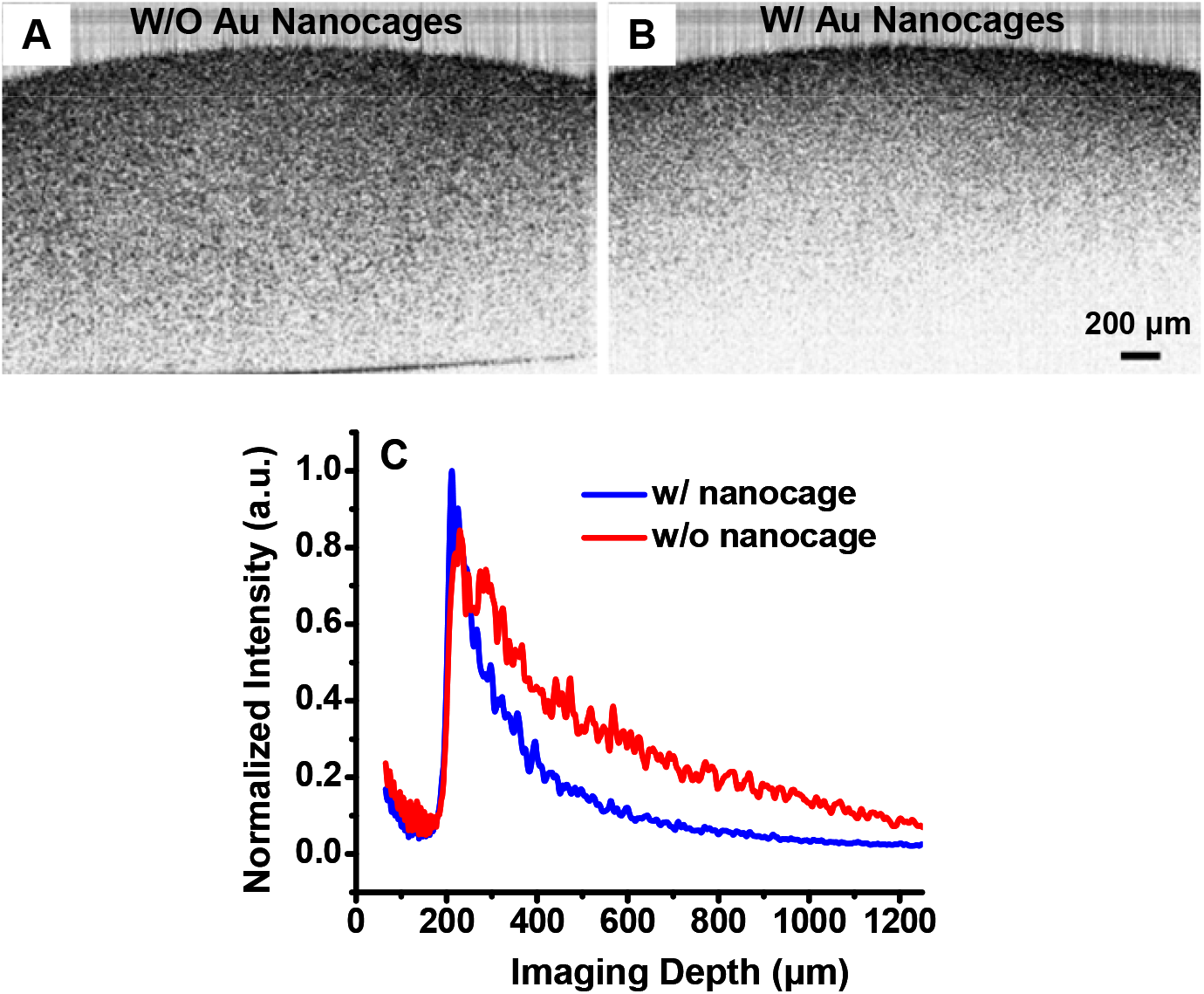
OCT images (obtained with a 1060 nm swept source) of a gelatin phantom embedded with TiO_2_ (1 mg/mL). (A and B) Intensity-based SS-OCT images of the phantom without and with gold nanocages, respectively. (C) SS-OCT intensity signals on a linear scale as a function of imaging depth. Note that the SS-OCT signal recorded from the portion of the phantom with gold nanocages decays faster than that from the portion without nanocages.

Figures 5A and 5B illustrate intensity-based SS-OCT images of the mouse tail tissue before and after the administration of gold nanocages, respectively. As in the phantom experiments, the presence of gold nanocages increased the total absorption of tissue. The nanocages were more or less uniformly distributed beneath the skin surface as they were locally injected.

**Figure 5.**
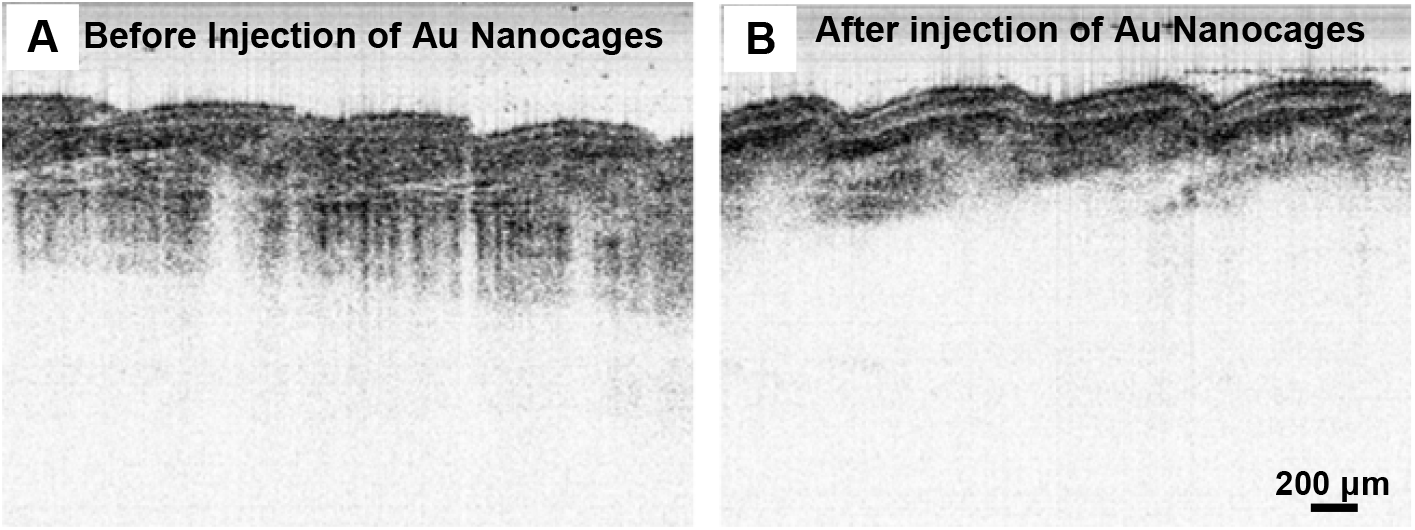
OCT images (with a 1060 nm swept source) of mouse tail tissue. (A) and (B) Intensity-based OCT images before and after the administration of nanocages (by local injection), respectively. The presence of gold nanocages increases the effective tissue absorption for intensity-based SS-OCT imaging.

## DISCUSSIONS

As demonstrated by the results shown in Figure 3, temperature and titration rate are critical parameters that control the reaction rate and allow for the tuning of the SPR peak wavelength to values above 1000 nm during the synthesis of gold nanocages. By lowering both the reaction temperature and titration rate compared to the conventional synthesis protocol for gold nanocages, we reduced the stress on the nanocage walls, thereby preventing collapse due to the stress corrosion cracking mechanism, as described in previous studies [17,18]. It is well-established that temperature has a significant effect on environmental cracking susceptibility, with crack velocity decreasing as the temperature drops, thus minimizing the apparent heat activation energies associated with the process [16]. Additionally, reducing the dealloying temperature significantly lowers the interfacial diffusivity of gold atoms, which leads to the formation of an ultrafine and stable nanoporous structure, thereby enhancing the chemical and physical properties of the nanocages [19,20]. Further reduction in the titration rate during the galvanic replacement reaction also helps reduce the potential stress on the walls of the nanocages, preventing collapse and promoting stability.

Under the reduced temperature and titration rate conditions, the SPR peak position could be precisely tuned to above 1000 nm by controlling the total volume of the HAuCl4 aqueous solution titrated into the reaction mixture, as shown in Figure 1. The SPR peak position of the gold nanocages is primarily determined by the wall thickness and porosity, which change during synthesis. As the porosity increases and the wall thickness decreases (as observed in Figure 2), the SPR peak shifts continuously to longer wavelengths. These observations are in agreement with our previous calculations based on discrete dipole approximation models for gold nanocages with an SPR peak around 800 nm (data not shown).

Integrating sphere measurements demonstrated that the gold nanocages had a large extinction cross-section (∼1.86 × 10^-15 m^2^) at 1060 nm, which is difficult to achieve with organic dyes. Compared with typical organic dyes such as Indocyanine Green (ICG), the extinction cross-section of the gold nanocages is approximately five orders of magnitude larger [21]. Furthermore, it was observed that the optical extinction at 1060 nm is dominated by absorption, as confirmed by the integrating sphere measurements. This strong absorption results in significant optical attenuation when gold nanocages are used as an absorbing imaging contrast agent. In highly scattering human tissues, such absorbing agents provide stronger contrast than scattering-based agents. This enhanced optical absorption contrast is clearly evident in the phantom and tissue OCT images (Figures 4B and 4C, and Figure 5B). Notably, this is the first demonstration of an OCT contrast agent using gold nanoparticles for imaging in the longer wavelength range (around 1060 nm). These results are consistent with our previous study, where gold nanocages were used as absorption-dominant contrast agents for OCT imaging at around 800 nm [9].

In summary, we presented a modified protocol for synthesizing gold nanocages that shifts their SPR peak towards longer wavelengths, specifically above 1000 nm. These nanocages represent a novel class of contrast agents for intensity-based OCT imaging at wavelengths beyond 1000 nm. Experiments with tissue phantoms and *in vivo* mouse tissues, using a 1060 nm SS-OCT system, demonstrated enhanced absorption contrast with the as-synthesized gold nanocages. Additionally, the strong absorption properties of the gold nanocages suggest potential applications for photothermal therapy, although this aspect is beyond the scope of the current study [22].

## Acknowledgements of Funding Supports

This work was supported in part by the National Institutes of Health (NIH) (R01 CA120480 and 1R01 EB007636) and the National Science Foundation (NSF) (IIP-0724231).

